# A new species of the genus *Microhyla* Tschudi, 1838 (Amphibia: Anura: Microhylidae) from eastern India with notes on Indian species

**DOI:** 10.1101/2021.08.07.455509

**Authors:** Somnath Bhakat, Soumendranath Bhakat

## Abstract

A new species of the genus *Microhyla*, *Microhyla bengalensis* sp. nov., described from West Bengal state, India. The new species is distinguished from its congeners by a combination of the following morphological characters: 1) Small in size (SVL= 16.2 mm. in male); 2) truncated snout in dorsal view; 3) head wider than long (HW: HL= 1.36); 4) canthus rostralis and tympanum are indistinct; 5) nostril placed on the dorsal side of the snout; 6) tibiotarsal articulation not reaching the eye; 7) fingers and toes without disc; 8) toe webbing basal; 9) thigh and foot length are equal and smaller than shank; 10) skin tuberculated on dorsum; 11) ‘teddy bear’ dark brown mark on dorsum; 12) an inverted ‘V’-shaped dark brown mark above the vent.

A comparative morphological data of all the 14 Indian species of *Microhyla* is also provided.

## Introduction

India is rich in amphibian diversity containing a significant number of newly discovered species. Exploration of amphibian species in India are mostly restricted to the diversity rich regions such as the Western Ghats in Peninsular India and the North east region (Biju et al., 2019). However, discoveries of new species from other regions of India particularly eastern part (including West Bengal) are scanty. Most new species discoveries are from Ranixalidae, Micrixalidae and Nyctibatrachidae families and a few from the cosmopoliton families like Microhylidae, Dicroglossidae and Bufonidae (Vineeth et al., 2018). The family Microhylidae consists of 13 subfamilies with worldwide distribution (Frost, 2018; Peloso et al., 2015). Of these, six genera and 26 species are found in India. The genus *Microhyla* Tschudi, 1838 is the most widely distributed group across Asia. Currently the genus is comprised of 52 recognised species (Li et al., 2019; Biju et al., 2019; Gorin et al.,2021) of which 13 species are known to occur in India, only four from north east region, one from Andaman and others from Western Ghats and southern states of India. Occurrence of *M. pulchra* and *M. butleri* in India is doubtful (Garg et al., 2019). The present paper describes a new species of the genus *Microhyla*, *Microhyla bengalensis* sp. nov. from a new distribution zone, West Bengal, an eastern state of India.

## Materials and methods

The discovery of the new species of *Microhyla* is purely an accident. On September 29, 2020 at 10.10 a. m. while clearing the leaf litter of a guava tree (*Psidium guajava* L.) in the courtyard, first author encountered the small frog. Initially, he ignored the specimen assuming it as an immature toad (*Duttaphrynus melanostictus* L.). Later, he observed it carefully and found the unique colouration and marking on the dorsum and then collected the specimen.

The specimen was euthenised, photographed and fixed in 4% formaldehyde for two days, finally washed and preserved in 70% ethyl alcohol.

Sex was determined by examining the gonads through a small ventral incision. Morphological measurements were taken to the nearest 0.1 mm. with the help of digital slide calipers, divider, centimeter scale and micrometer. The specimen was also examined under a streomicroscope to study the morphometric and meristic characters. The following abbreviations were used in the text: SVL (snout-vent length), HW (head width, at the angle of jaws), HL (head length, from rear of the mandible to the tip of the snout), SL (snout length, from anterior orbital border to the tip of the snout), EL (eye length, horizontal distance between the orbital borders), EN ( distance from the front of the eye to the nostril), NS (distance from the nostril to the tip of the snout), IN (internarial distance), IUE (inter upper eye lid width, shortest distance between the upper eye lid), UEW (maximum upper eye lid width), IFE (internal front of the eyes, shortest distance between the anterior orbital borders), IBE (internal back of the eyes, shortest distance between the posterior orbital borders), MN (distance from rear of the mandible to nostril), MFE (distance from rear of the mandible to the anterior orbital border of the eye), MBE (distance from rear of the mandible to the posterior orbital border of the eye), FAL (forearm length, from the flexed elbow to the base of the outer palmer tubercle), HAL (hand length, from the base of the outer palmer tubercle to the tip of the third finger), THL (thigh length, from vent to knee), SHL (shank length, from knee to heel), FOL (foot length, from the base of the inner metatarsal tubercle to the tip of the fourth toe), TFOL (distance from the heel to the tip of the fourth toe), IMT (length of inner metatarsal tubercle), OMT (length of the outer metatarsal tubercle). Digit numbers are represented by roman numerals, I – V. All morphometric measurements in the text are in mm. Size and webbing of the species are categorized following Garg et al. (2018) and Garg and Biju (2017). Toe webbing and subarticular tubercle formula were in accordance with those of Savage (1975).

Comparisons were made with the data taken from literature (Vogt, 1911; Inger, 1989; Pillai, 1977; Dutta and Ray, 2000; Bain and Nguyen, 2004; Matsui et al. 2013; Hasan et al, 2014; Poyarkov Jr. et al., 2014; Howlader et al., 2015; Seshadri et al., 2016a, 2016b; Khatiwada et al., 2017; Vineeth et al., 2018; Biju et al., 2019; Garg et al., 2019; and Li et al., 2019). Paired t-test was done to evaluate the separation of the morphometric variables between the new species and other 13 existing Indian species of *Microhyla*.

Principal component analysis (PCA) and construction of heat map was performed on the multivariate data of 14 species of *Microhyla*, using Clust Vis: https://biit.cs.ut.ee/clustvis/ (Metsalu and Vilo, 2015). PCA is a technique to represent high dimensional data and using the dependent variables in a more tractable, lower dimensional form. Plotting PC1 vs. PC2 approximate the distance between points which form separate clusters.

Heat map uses colour coding to visualize multivariate data matrix. The phenogram integrated with heat map shows how corresponding variables (species or character) are separated better than others.

## Results

### Systematic position

Amphibia Linnaeus, 1758

Anura Fisher von Waldheim, 1813

Microhylidae Gunther, 1858

Microhylinae Gunther, 1858

*Microhyla* Tschudi, 1838

*Microhyla bengalensis* sp. nov. (Fig. 1 and 2, Table I).

**Fig. 1.**
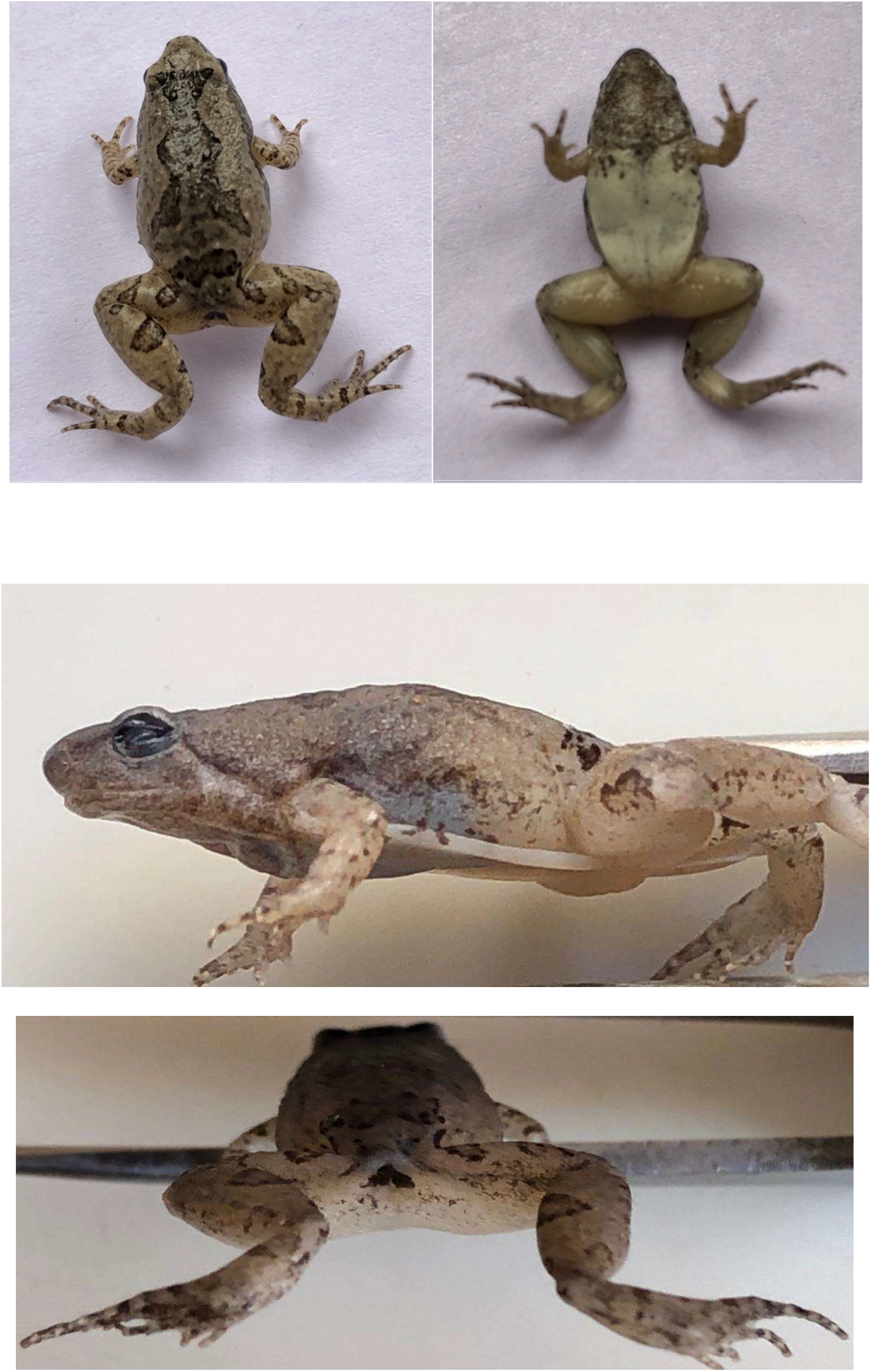
*Microhyla bengalensis* sp. nov. Upper: Dorsal and ventral view, Middle: Lateral view, Lower: Showing vent with dark brown mark.

**Fig. 2.**
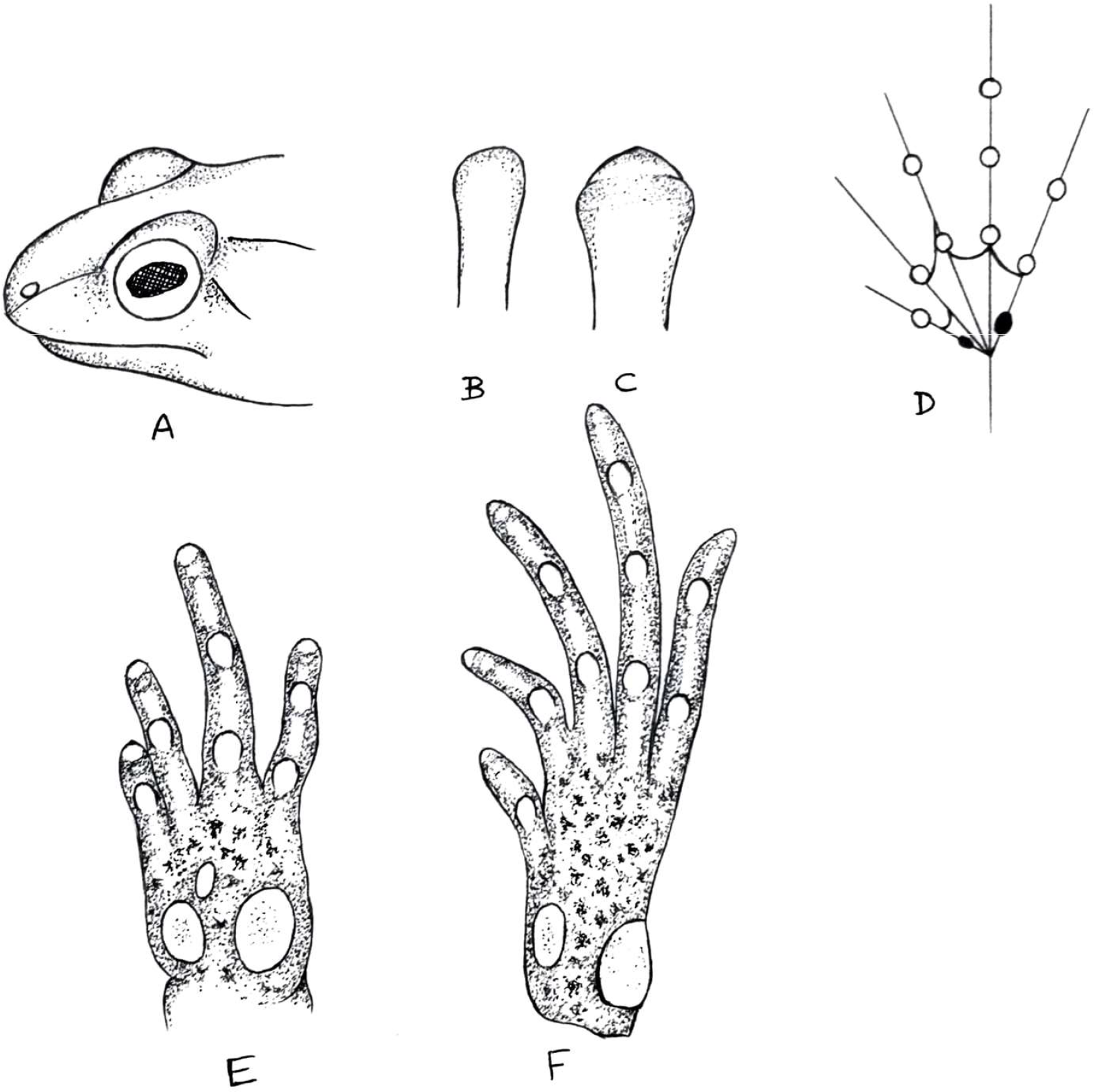
*Microhyla bengalensis* sp. nov. A. nostril placed dorsally, B. third finger tip, C. fourth toe tip, D. webbing of foot, E. ventral view of hand, F. ventral view of foot.

**Table 1.** Morphometric measurements of *Microhyla bengalensis* sp. nov.

| Sl. No. | Paramaters | Values | % SVL | Sl. No. | Parameters | Values | % SVL |
| --- | --- | --- | --- | --- | --- | --- | --- |
| 1. | SVL | 16.2 |  | 17. | HAL | 3.9 | 24.07 |
| 2. | HW | 6.0 | 37.04 | 18. | THL | 7.1 | 43.83 |
| 3. | HL | 4.4 | 27.16 | 19. | SHL | 8.3 | 51.23 |
| 4. | SL | 1.6 | 9.88 | 20. | FOL | 7.2 | 44.44 |
| 5. | EL | 2.0 | 12.35 | 21. | TFOL | 11.2 | 69.14 |
| 6. | EN | 0.8 | 4.94 | 22. | FLI | 1.0 | 6.17 |
| 7. | NS | 0.4 | 2.47 | 23. | FLII | 1.3 | 8.02 |
| 8. | IN | 1.8 | 11.11 | 24. | FLIII | 3.0 | 18.52 |
| 9. | IUE | 1.5 | 9.26 | 25. | FLIV | 2.4 | 14.81 |
| 10. | UEW | 1.2 | 7.41 | 26. | TLI | 1.4 | 8.64 |
| 11. | IFE | 3.3 | 20.37 | 27. | TLII | 2.5 | 15.43 |
| 12. | IBE | 4.4 | 27.16 | 28. | TLIII | 4.0 | 24.69 |
| 13. | MN | 4.6 | 28.40 | 29. | TLIV | 5.4 | 33.33 |
| 14. | MFE | 3.5 | 21.60 | 30. | TLV | 3.9 | 24.07 |
| 15. | MBE | 2.0 | 12.35 | 31. | IMT | 0.5 | 3.09 |
| 16. | FAL | 3.6 | 22.22 | 32. | OMT | 0.9 | 5.55 |

### Etymology

The species name is derived from the name of the type locality ‘Bengal’ (West Bengal state) where the holotype was collected.

### Holotype

Zoological Museum, Dept. of Zoology, Rampurhat College, Rampurhat-731224, Dist. Birbhum, W. B. India; an adult male collected from leaf litters of a guava tree in Vivekanandapally, Suri (87°32′00′′E, 23°55′00′′N), Birbhum district, West Bengal, India.

### Diagnosis

The new species was assigned to the genus *Microhyla* based on the set of following characters as described by Parker (1934), Inger (1989), Matsui et al. (2013), Poyarkov et al. (2014), Wijayathilaka et al. (2016), Seshadri et al. (2016a), Khatiwada (2017) and Garg et al. (2019): small sized adult, narrow head wider than long, snout less than twice the diameter of the eye, indistinct canthus rostralis, tympanum hidden by skin, eyes with circular pupil, presence of supratympanic fold, vomerine teeth absent, tongue entire without papilla, distinct inner and outer metatarsal tubercle, absence of webbing between fingers and absence of supernumery tubercles on fingers and toes.

The new species is diagnosed by the following set of characters: small sized adult (male, SVL: 16.2 mm.), snout truncated in dorsal but triangular in ventral view, canthus rostralis and tympanum indistinct; snout protrudes beyond mouth in ventral view; vomerine teeth absent; tongue tape like with flat tip, margin smooth and without lingual papillae; nostril placed dorsally; a lateral dermal fold from behind eye to shoulder; no dorsal median line; first finger is longer than half the length of the second finger; fingers without webbing; toes with basal webbing; tibiotarsal articulation not reaching eye; head triangular, wider than long; palmer tubercles well developed; inner metatarsal tubercle is ovoid; outer metatarsal tubercle large and shovel shaped; skin tuberculated on dorsum; dark brownish pigmented throat; vent with an inverted ‘V’ shaped dark brown mark; posterior of thigh white and granulated; dorsum with dark brown “teddy bear” shaped marking from inter-orbital line to sacral region. Limbs with brown hollow marking, fingers and toes with brown cross bars.

### Description of holotype

Small sized adult male (SVL= 16.2 mm.). Snout truncated in dorsal, triangular in ventral and rounded in lateral view (Fig.1); head wider than long (HW: HL= 1.36); upper jaw distinctly protrudes in lateral view, junction of upper and lower jaw extended beyond middle of eye length. Snout length (SL= 1.6) is shorter than horizontal diameter of the eye (EL= 2.0); canthus rostralis indistinct; interorbital space is more or less flat and wider (IUE= 1.5) than upper eyelid width (UEW= 1.2) and shorter than internarial distance (IN= 1.8). The distance between posterior margin of eyes (IBE= 4.4) 1.33 times that of anterior margins (IFE= 3.3). Nostrils are small and rounded without flap, placed on the dorsal side of snout (Fig. 2). The distance between nostril to eye is two times that of nostril to snout (EN= 0.8, NS= 0.4); tympanum indistinct; distinct supratympanic fold extending from posterior corner of eye to the shoulder (Fig. 1); vomerine teeth absent; tongue long tape like without papillae. Eye is small and 1.33 times of interorbital width, pupil rounded. Fore arm length (FAL= 3.6) is shorter than hand length (HAL= 3.9); fingers short, relative length of fingers: I< II< IV< III (FLI= 1, FLII= 1.3, FLIII= 3, FLIV= 2.4). Disc on finger tips and webbing between fingers are absent, tips slightly wider compared to finger widths; palmer tubercle well developed and with a small accessory palmer tubercle, thenar tubercle is large, ovoid and laterally placed; subarticular tubercles very distinct, ovoid and protruded, finger subarticular tubercle formula: 1: 1: 2: 2 (Fig. 2). Nuptial pad is absent. Hind limb is about 1.64 times SVL (HLL= 26.6, SVL= 16.2). Tibiotarsal articulation of adpressed hind limb not reaching the eye when leg stretched forward, heels overlap when thighs are positioned at right angles to the body. Shank is longer than both thigh and foot length which are almost equal in (SHL= 8.3, THL= 7.1, FOL= 7.2). Heel to tip of fourth toe (TFOL= 11.2) about 4.67 times longer than fourth toe length (TLIV= 2.4); relative length of toes: I< II< V< III< IV (TLI= 1.4, TLII= 2.5, TLIII= 4.0, TLIV= 5.4, TLV= 3.9), toe tips rounded without disc and longitudinal grooves; toe webbing reduced: I2^1/2^ – 2^1/2^ II2 – 3^−^III3^+^ −4^+^IV4^+^ −3^+^V (Fig. 2). Subarticular tubercle distinct, oval in shape and protruding; toe subarticular tubercles formula: 1: 1: 2: 3: 2. Inner metatarsal tubercle is elongated (IMT= 0.5) and almost half of outer metatarsal tubercle (OMT= 0.9) which is large and shovel-shape, supernumery tubercles absent (Fig. 2).

### Skin texture and colour

Dorsal skin surface rough with tiny tubercles, more on the posterior end, lateral side of the body is granulated, cloacal region and posterior of thigh granulated. Skin on ventral side is smooth, throat shagreened.

Dorsum yellowish brown, with a dark brown “teddy bear” shape marking extending from interorbital line to sacral region, at the posterior end of the mark two lateral slender lobes are extended. At the caudal region, a distinct dumbel shaped dark brown mark is present. Thin dark brown line arises from anterior corner of eye along canthus rostralis toward nostril. Flanks are with an irregular line of brown spots starting above the shoulder and terminating well short of groin. Upper portion of vent bears an inverted ‘V’ mark, dark brown in colour. Posterior of thigh is white in colour. A white band of muscle fold extending from fore limb to thigh on both sides of the body; fingers and toes with narrow dark brown bars in dorsal view, palm and sole with dark brown pigments, subarticular tubercles are crystal white. Abdomen is creamy white, throat and chest with dark brown pigments. Iris is black in colour. Fore arm with brown ring and hand with brown bars and spots. Each thigh with two distinct spots, one circular deep brown hollow spot at proximal region and another inverted ‘,’ shape brown hollow mark in the mid region. Shank with a ‘>’shape hollow dark brown mark in the middle and two round hollow brown marks one at the knee and other at the heel end; foot have dark brown bars (Fig. 1).

### Comparison

Garg et al. (2019) categorized the male *Microhyla* species on the basis of snout-vent length (SVL) as: small (SVL= 13-20 mm), medium (21-30 mm.) and large (SVL > 31 mm.). On this basis the new species, *Microhyla bengalensis* (SVL= 16.2 mm.) belongs to the first category i. e. small. Biju et al. (2019) listed the snout-vent length (SVL) of all the 50 recognised species of *Microhyla* (based on the studies of Garg et al. 2018a, Nguyen et al. 2019 and Li et al. 2019). Since the present species is a congener of small microhylids by size ranges, a detailed comparison with 10 Indian species viz. *M. chakrapanii*, *M. darreli*, *M. heymonsi*, *M. kodial*, *M. laterite*, *M. mukhlesuri*, *M. mymensinghensis,*. *M. nilphamariensis*, *M. ornata*, *M. sholigari* and 24 non-Indian species viz. *M. annamensis*, *M. annectens, M. arboricola*, *M. beilunensis*, *M. borneensis*, *M. fanjingshanensis*, *M. gadjahmadai*, *M. irrawaddy*, *M. karunaratnei*, *M. maculifera*, *M. malang*, *M. mantheyi*, *M. marmorata*, *M. minuta*, *M. mixtura*, *M. orientalis*, *M. palmipes*, *M. perparva*, *M. petrigena*, *M. pineticola*, *M. pulchella*, *M. pulverata*, *M. taraiensis* and *M. zeylanica* is provided. Among non-Indian species except five species: *M. beilunensis*, *M. fanjingshanensis*, *M. irrawaddy*, *M. maculifera* and *M. taraiensis*, all have distinct disc on toes and fingers. The present species differs from those 19 species by lacking of disc on toes and fingers (vs. presence of disc on toes and fingers). The new species differs from *M. beilunensis*, *M. fanjingshanensis* and *M. taraiensis* by the presence of truncated snout (vs. rounded snout in other three species). It differs from *M. maculifera* by the presence of two palmer tubercles and rudimentary webbing on toes (vs. one palmer tubercle and webbed toes in *M. maculifera*). Further, *M. maculifera* is relatively smaller in size (SVL= 12.0 – 13.3 vs. SVL= 16.2 in *M. bengalensis*). It differs from *M. irrawaddy* by greater EL/SL ratio (1.25 vs. 0.88), smaller thigh length ratio (THL: SVL= 0.44 vs. 0.51), proportionate length of finger and toe (in *M. bengalensis* finger: I< II< IV< III vs. I< II= IV< III in *M. irrawaddy*, toe: I< II< V< III< IV in *M. bengalensis* vs. I< V< II< III< IV in *M. irrawaddy*), larger OMT (vs. smaller). Further, tibiotarsal articulation in *M. irrawaddy* reaches the eye level while in *M. bengalensis* not reaches the eye level. *M. bengalensis* differs from *M. taraiensis* by larger OMT (OMT: IMT= 1.8 vs. 0.5 in *M. taraiensis*); smaller HL/HW ratio (0.73 vs. 0.83 – 0.96); larger eye (EL: SVL= 12.35 vs. 7.19); larger SHL: SVL ratio (51.23 vs. 31.13).

In all 10 small-sized Indian species of *Microhyla*, snout is acutely pointed or rounded except *M. nilphamariensis* in which snout is truncated. The new species is very similar to the aforementioned species in having a truncated snout. Further, *M. bengalensis* differs from other six species of small-sized Indian species (*M. chakrapanii*, *M. darreli*, *M. heymonsi*, *M. kodial*, *M. laterite* and *M. sholagari*) lacking digital disc (vs. present in all six species).

*M. bengalensis* differs from *M. chakrapanii* by smaller SL (9.88 vs. 11.36 %SVL) and IUE (9.26 vs. 13.64%SVL) and longer hind limb (164.19 vs. 145.45%SVL), Further, UEW is 23% wider in *M. chakrapanii* that of *M. bengalensis* and tibiotarsal articulation in *M. chakrapanii* just reaches eye level (vs. not reaches eye level in *M. bengalensis*). The new species differs from *M. darreli* in having higher HW: HL ratio (1.36 vs. 1.05), shorter snout (9.88 vs. 12.58 %SVL), smaller NS (2.47 vs. 4.64 %SVL), and larger eye length (12.35 vs. 8.61 %SVL). Both IN and UEW in *M. bengalensis* is 39% greater than those of *M. darreli*. All the units of hind limb (THL, SHL, FOL and TFOL) are proportionately greater in *M. darreli* (48.34 vs. 43.83, 54.97 vs. 51.23, 52.98 vs. 44.44 and 78.15 vs. 69.14 in %SVL respectively). A narrow mid-dorsal skin fold or line extending from tip of snout to vent is present in *M. darreli* (vs. absent in *M. bengalensis*). Further, relative lengths of fingers and toes are also different in these two species. The present species differs from *M. heymonsi* by greater HW: HL ratio (1.36 vs. 1.08) and absence of ‘()’ shaped marking on the dorsum (vs. present). SL and NS in *M. heymonsi* are greater than those of *M. bengalensis* (12.38 vs. 9.88 %SVL and 4.46 vs. 2.47 %SVL respectively) while reverse are true in case of EL and IN (8.91 vs. 12.35 %SVL and 8.91 vs. 11.11 %SVL respectively). FAL in *M. bengalensis* is 21% greater that of *M. heymonsi* but THL and SHL are shorter (25% and 19% respectively) those of *M. heymonsi*. Further, ratios of FOL: SVL and TFOL: SVL in *M. bengalensis* are smaller than those of *M. heymonsi* (0.44 vs. 0.62 and 0.69 vs. 0.88 respectively). *M. bengalensis* differs from *M. kodial* by greater HW: HL ratio (1.36 vs. 1.12), shorter snout length (9.88 vs. 12.59 %SVL), smaller EN: SVL ratio (4.94 vs. 7.69), longer eye length (12.35 vs. 9.32 %SVL). In *M. kodial*, SHL and FOL are almost equal in length while in *M. bengalensis*, SHL is 15% longer than FOL. Further, MN, MBE and MFE are proportionately more in *M. bengalensis* compared to those of *M. kodial* (28.40 vs. 18.53, 12.35 vs. 9.56 and 21.60 vs. 14.80 in %SVL respectively). The present species differs from *M. laterite* by greater HW: HL ratio (1.36 vs. 1.20), shorter snout length (9.88 vs. 12.35 %SVL), greater IN: SVL ratio (11.11 vs. 8.81), smaller EN: SVL ratio (4.94 vs. 7.35). NS of *M. laterite* is two times that of *M. bengalensis* (5.04 vs. 2.47 %SVL). MN, MBE and MFE are proportionately more in *M. bengalensis* than those of *M. laterite* (28.40 vs. 20.41, 12.35 vs. 10.93 and 21.60 vs. 15.33 in %SVL respectively). In *M. laterite*, SHL and FOL are almost equal in length (ca. 50 %SVL) while SHL (51.23 %SVL) is longer than THL (43.83 %SVL) in *M. bengalensis*. Further, OMT: IMT ratio is more in *M. bengalensis* (1.8 vs. 0.63 in *M. laterite*). *M. bengalensis* differs from *M. mukhlesuri* by greater ratios of HW: HL (1.36 vs. 0.85) and IN: SVL (11.11 vs. 8.38); smaller ratios of NS: SVL (2.47 vs. 3.91), EN: SVL (4.94 vs. 9.50), IUE: SVL (9.26 vs. 12.29). Width of upper eye lid in *M. bengalensis* is 20 % more compared to that of *M. mukhlesuri.* The new species differs from *M. mukhelsuri* by its tibiotarsal articulation reaching until the eye (vs. reaching eye to the tip of the snout) and finger formula (I< II< IV< III vs. I< IV< II< III in *M. mukhlesuri*). It differs from *M. mymensinghensis* by greater ratios of HW: HL (1.36 vs. 1.22), IN: SVL (11.11vs. 9.86) and smaller ratios of NS: SVL (2.47 vs. 6.10), IUE: SVL (9.26 vs. 13.62) and UEW: SVL (7.41 vs. 9.86). Distance between eye to nostril is more than double in *M. mymensinghensis* that of *M. bengalensis* (10.80 vs. 4.94 %SVL). Further, thigh and shank length is almost equal in *M. mymensinghensis* while thigh length is smaller than shank length in *M. bengalensis* (THL: SHL= 0.85). The new species differs from *M. mymensinghensis* by skin texture also (tuberculated vs. smooth in *M. mymensinghensis*). The new species differs from *M. nilphamariensis* by longer head (27 vs. 21 %SVL), distance from back of mandible to back of eye 45% of head length (vs. 15% of head length in *M. nilphamariensis*). Further, it differs from *M. nilphamariensis* by greater UEW: EL ratio (0.60 vs. 0.44), and smaller ratios of EI: HL (0.45 vs. 0.52), EN: NS (2.0 vs. 5.95), IUE: IN (0.83 vs. 2.03) and SL: HL (0.36 vs. 0.47). OMT in *M. nilphamariensis* is minute and indistinct while it is large, distinct and greater than IMT. *M. bengalensis* differs from *M. sholigari* by greater ratios of HW: HL (1.36 vs. 1.24), EL: SVL (12.35 vs. 10.76), IN: SVL (11.11 vs. 9.07) and IFE: SVL (20.37 vs. 16.15) and smaller ratio of SL: SVL (9.88 vs. 14.21). Snout in *M. sholigari* 1.32 times longer than eye length (vs. shorter than eye length in *M. bengalensis*, SL: EL= 0.8) and IUE is 1.68 times than UEW (vs. 1.25 times in *M. bengalensis*). Hind limb of *M. bengalensis* is 1.64 times longer than SVL (vs. 2.0 times in *M. sholigari*). Further, units of hind limb (THL, SHL, TFOL) are also smaller in *M. bengalensis* those of *M. sholigari* (43.83 vs. 54.13, 51.23 vs. 59.57 and 69.14 vs. 85.73 %SVL respectively). MN, MBE and MFE in *M. bengalensis* are greater in comparison with those of *M. sholigari* (28.40 vs. 21.59, 12.35 vs. 8.01 and 21.60 vs. 15.46 %SVL respectively). OMT in the present species is larger than IMT while reverse is true in *M. sholigari* (OMT: IMT= 1.8 vs. 0.27 in *M. sholigari*). Like most of the small sized microhylid, supernumery tubercles are absent in *M. bengalensis* (vs. present in *M. sholigari*). The new species differs from *M. ornata* by greater ratios of HW: HL (1.36 vs. 1.25), EL: SVL (12.35 vs. 8.30), UEW: SVL (7.41 vs. 5.24) and smaller ratios of EN: SVL (4.94 vs. 5.68), NS: SVL (2.47 vs. 3.49) and IUE: SVL (9.26 vs. 10.480. Snout of *M. bengalensis* is ca. 20% longer that of *M. ornata*. Units of fore limb (FAL and HAL) in *M. bengalensis* are more in length than those of *M. ornata* (22.22 vs. 16.16% SVL and 24.07 vs. 21.40% SVL respectively). Hind limb in new species is more than 10% greater in length that of *M. ornata*. Further, the present species differs from *M. ornata* by the presence of OMT (vs. absent in *M. ornata*) and its tibiotarsal articulation reaching until the eye (vs. level of shoulder in *M. ornata*). The result of the paired t-test indicates that except four species viz. *M. berdmorei*, *M. chakrapanii*, *M. heymonsi* and *M. eos*, all nine species are significantly different from the present species on the basis 20 or more morphological characters (Table 2). A comparison of characters of all known 14 Indian species of *Microhyla* is given in Table 3.

**Table 2.**
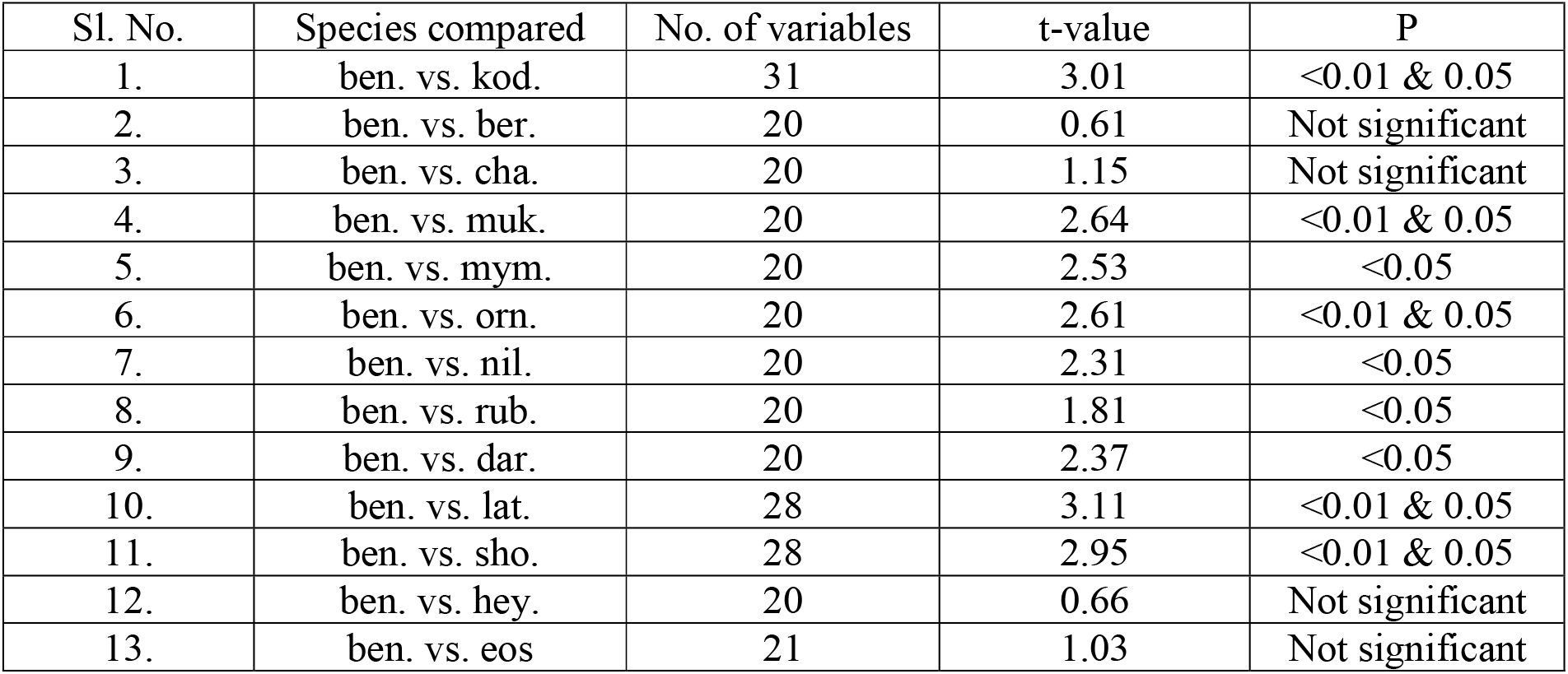
Statistics obtained from pair wise comparison using measurements of 14 Indian species of *Microhyla.* Abbreviations: ben.= *M. bengalensis*, kod.= *M. kodial,* ber.= *M. berdmorei*, cha= *M. chakrapanii*, muk.= *M. mukhlesuri*, mym.= *M. mymensinghensis*, orn.= *M. ornata*, nil= *M. nilphamariensis*, rub.= *M. rubra*, dar.= *M. darreli*, lat.= *M. laterite*, sho.= *M. sholigari*, hey.= *M. heymonsi*, eos.= *M. eos*.

| Sl. No. | Species compared | No. of variables | t-value | P |
| --- | --- | --- | --- | --- |
| 1. | ben. vs. kod. | 31 | 3.01 | <0.01 & 0.05 |
| 2. | ben. vs. ber. | 20 | 0.61 | Not significant |
| 3. | ben. vs. cha. | 20 | 1.15 | Not significant |
| 4. | ben. vs. muk. | 20 | 2.64 | <0.01 & 0.05 |
| 5. | ben. vs. mym. | 20 | 2.53 | <0.05 |
| 6. | ben. vs. orn. | 20 | 2.61 | <0.01 & 0.05 |
| 7. | ben. vs. nil. | 20 | 2.31 | <0.05 |
| 8. | ben. vs. rub. | 20 | 1.81 | <0.05 |
| 9. | ben. vs. dar. | 20 | 2.37 | <0.05 |
| 10. | ben. vs. lat. | 28 | 3.11 | <0.01 & 0.05 |
| 11. | ben. vs. sho. | 28 | 2.95 | <0.01 & 0.05 |
| 12. | ben. vs. hey. | 20 | 0.66 | Not significant |
| 13. | ben. vs. eos | 21 | 1.03 | Not significant |

**Table 3.**
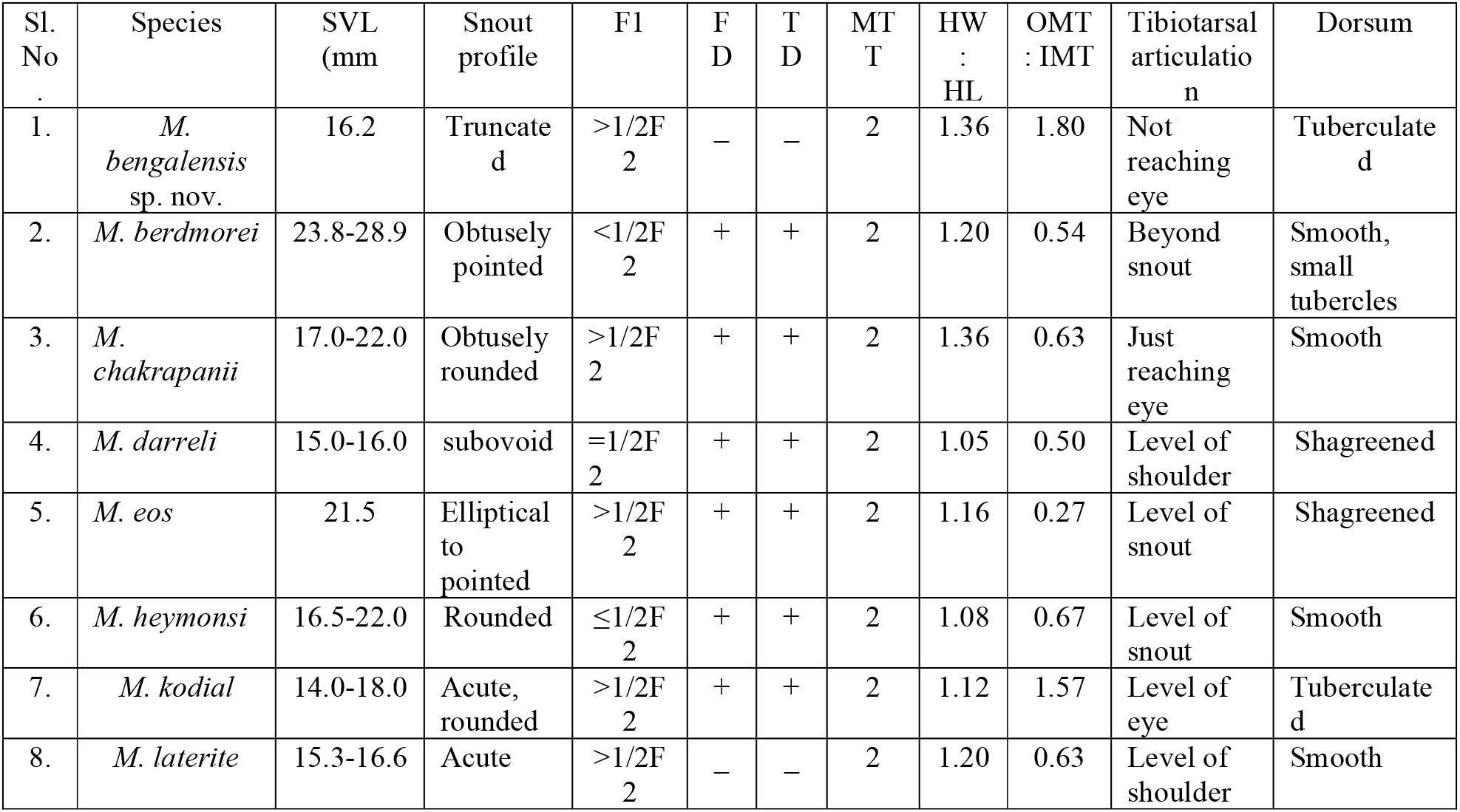
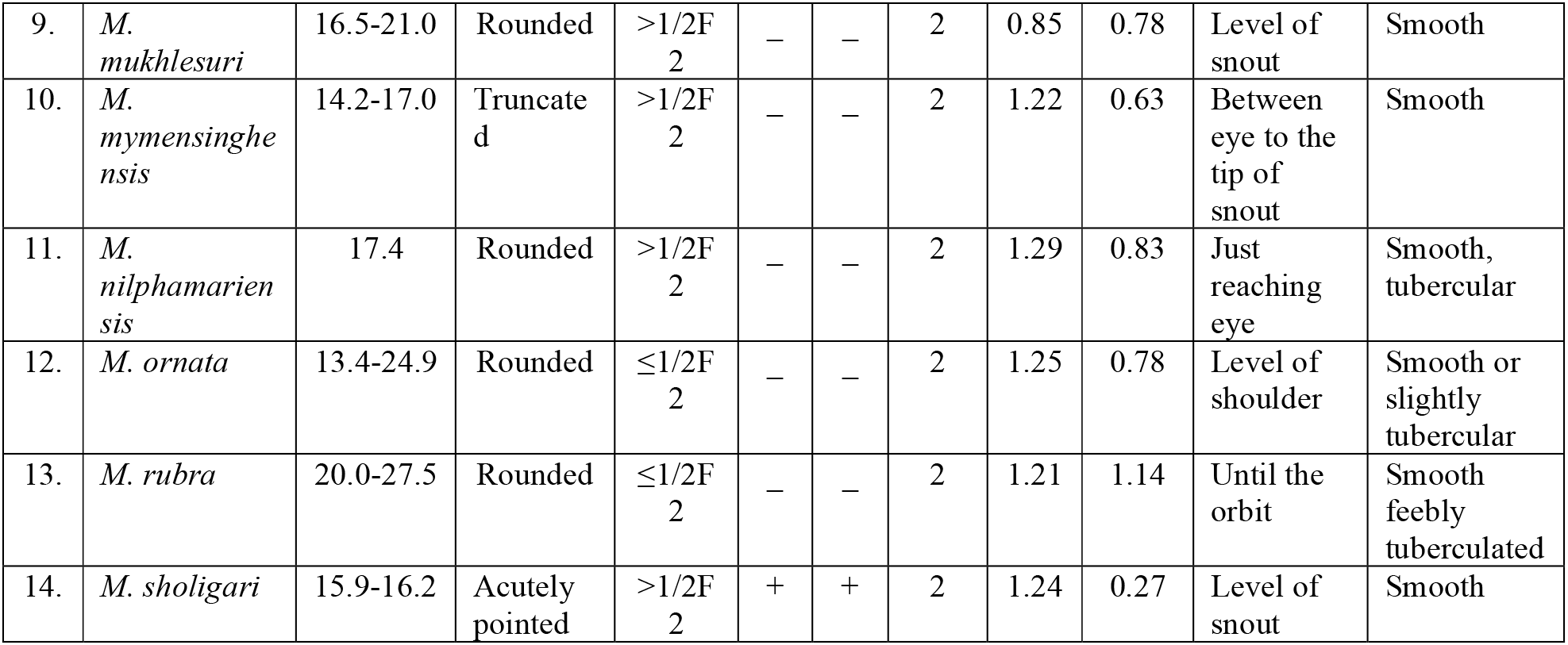
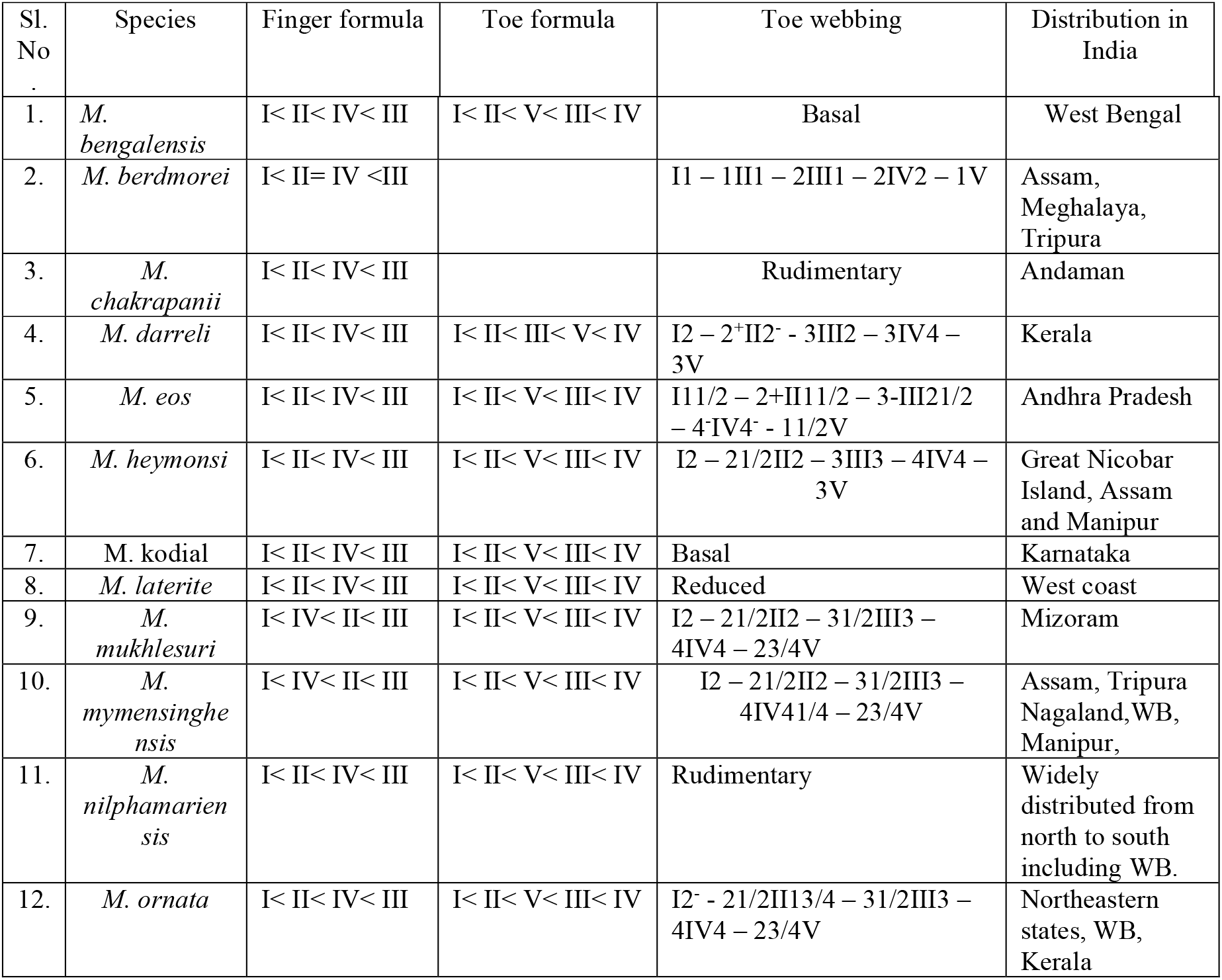
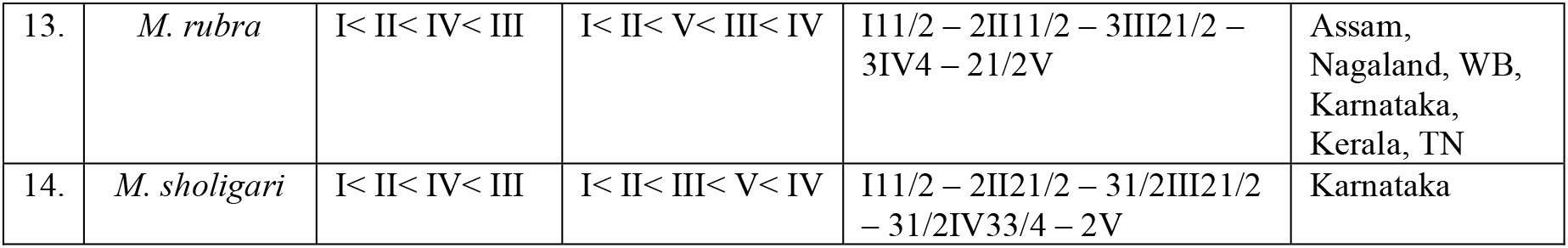
Comparative table of different morphological character and distribution of 14 Indian species of *Microhyla*. Abbreviations: F1= First finger, F2= Second finger, FD= Finger disc, TD= Toe disc, MTT= Metatarsal tubercle, HW= Head width, HL= Head length, OMT= Outer metatarsal tubercle, IMT= Inner metatarsal tubercle.

PCA of multivariate data of 14 species of *Microhyla* is presented in Fig. 4. PC1 and PC2 capture 57.8% variance in our data. Projection of PCs along first two components indicate that three species, *M. chakrapani*, *M. laterite* and *M. darreli* are closed to each other. PCs clearly shows that based on morphological features, *M. bengalensis* is a distinct species and separated from all other Indian species of *Microhyla.*

**Fig. 3.**
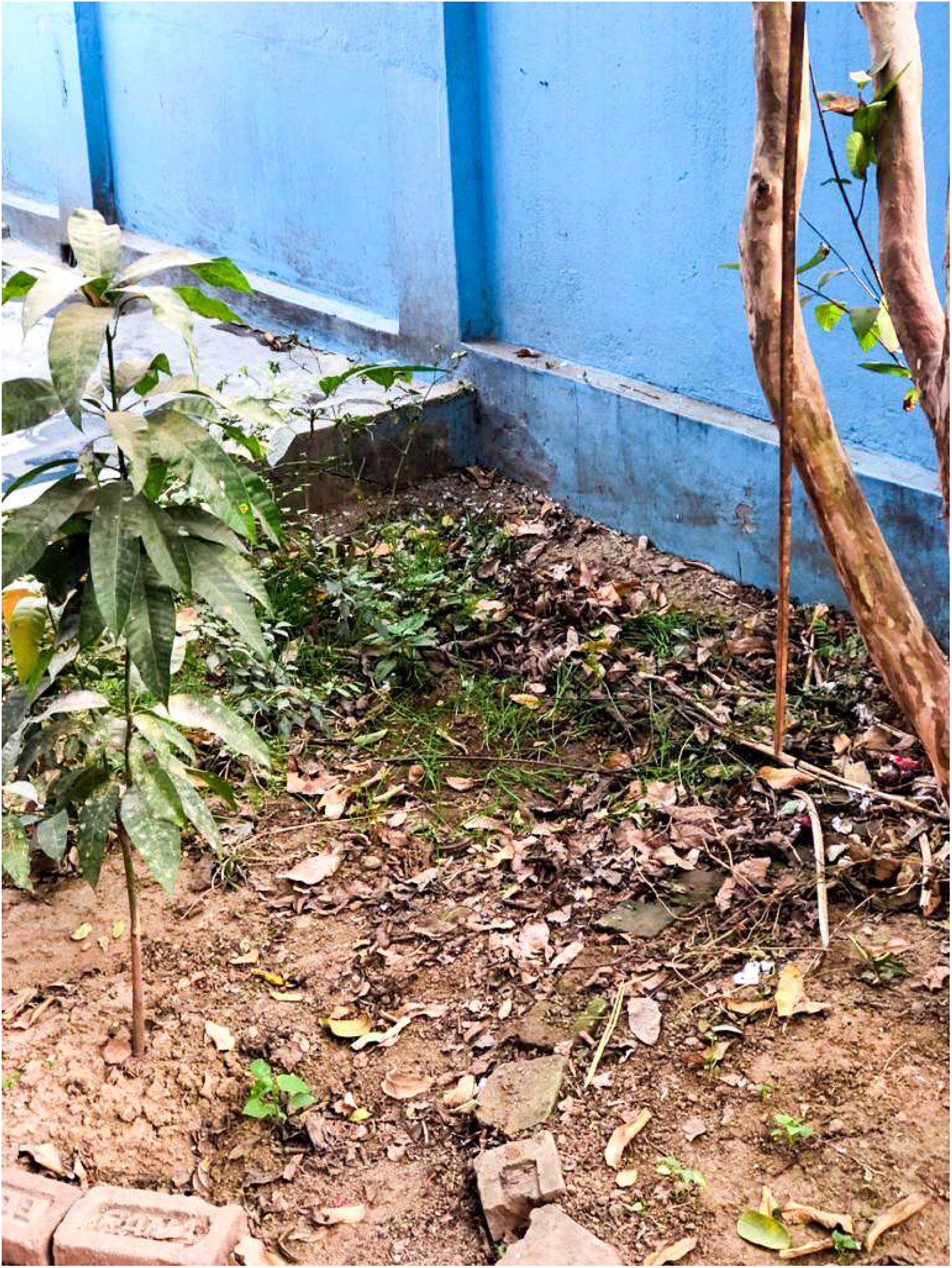
Type locality of *Microhyla bengalensis* sp. nov.

**Fig. 4.**
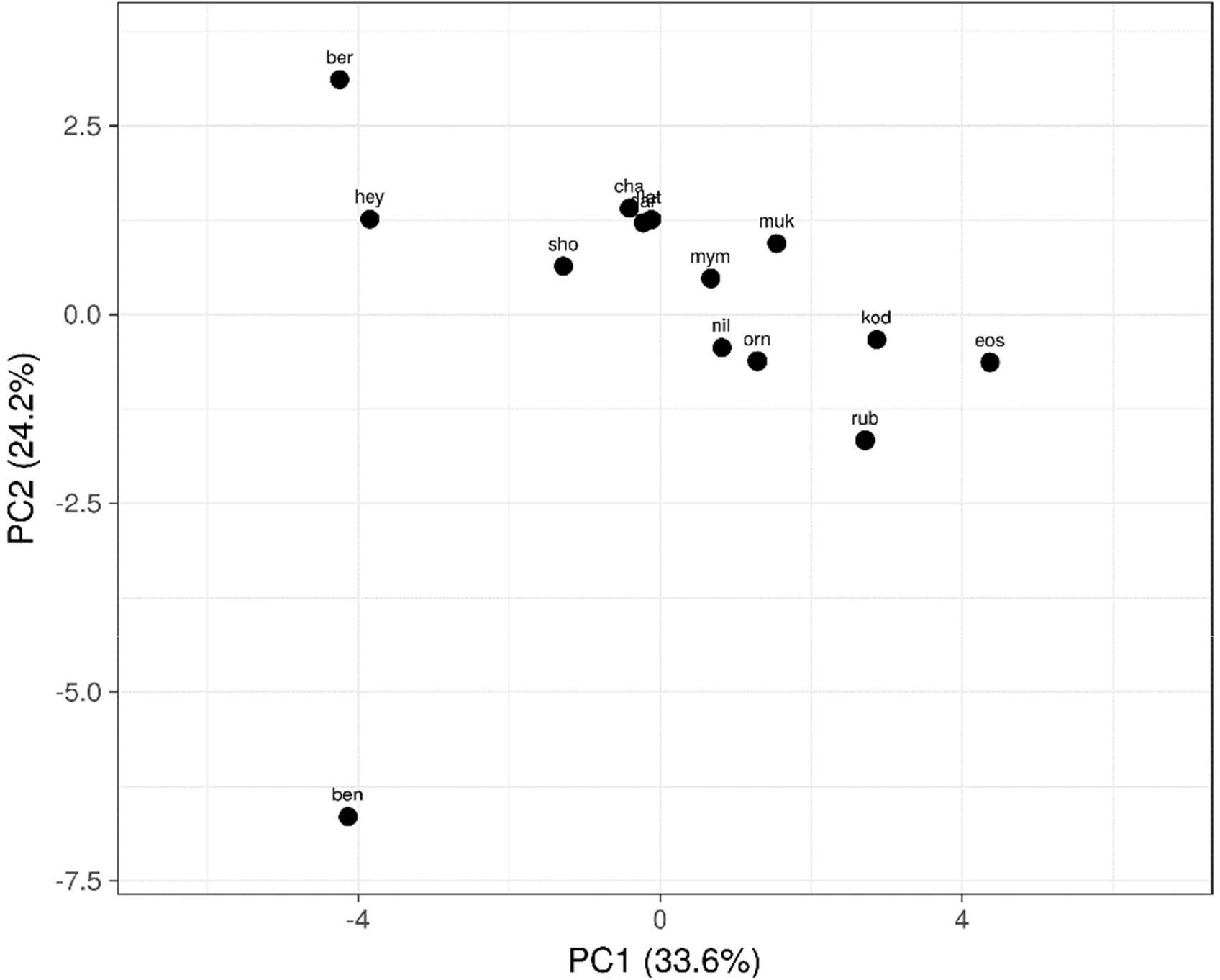
Principal Component Analysis (PCA) of multivariate data of 14 species of *Microhyla* (for abbreviations – Table 2).

Heat map integrated with phenogram is presented in Fig. 5. Phenogram shows that for some morphological features (FI, FIII, EL, IN, HW, FAL), *M bengalensis* is clearly separated from other 13 species of *Microhyla*. Moreover, species phenogram indicate that *M. bengalensis* acts as linker in between two major groups of *Microhyla* (one consists of six and another with seven species).

**Fig. 5.**
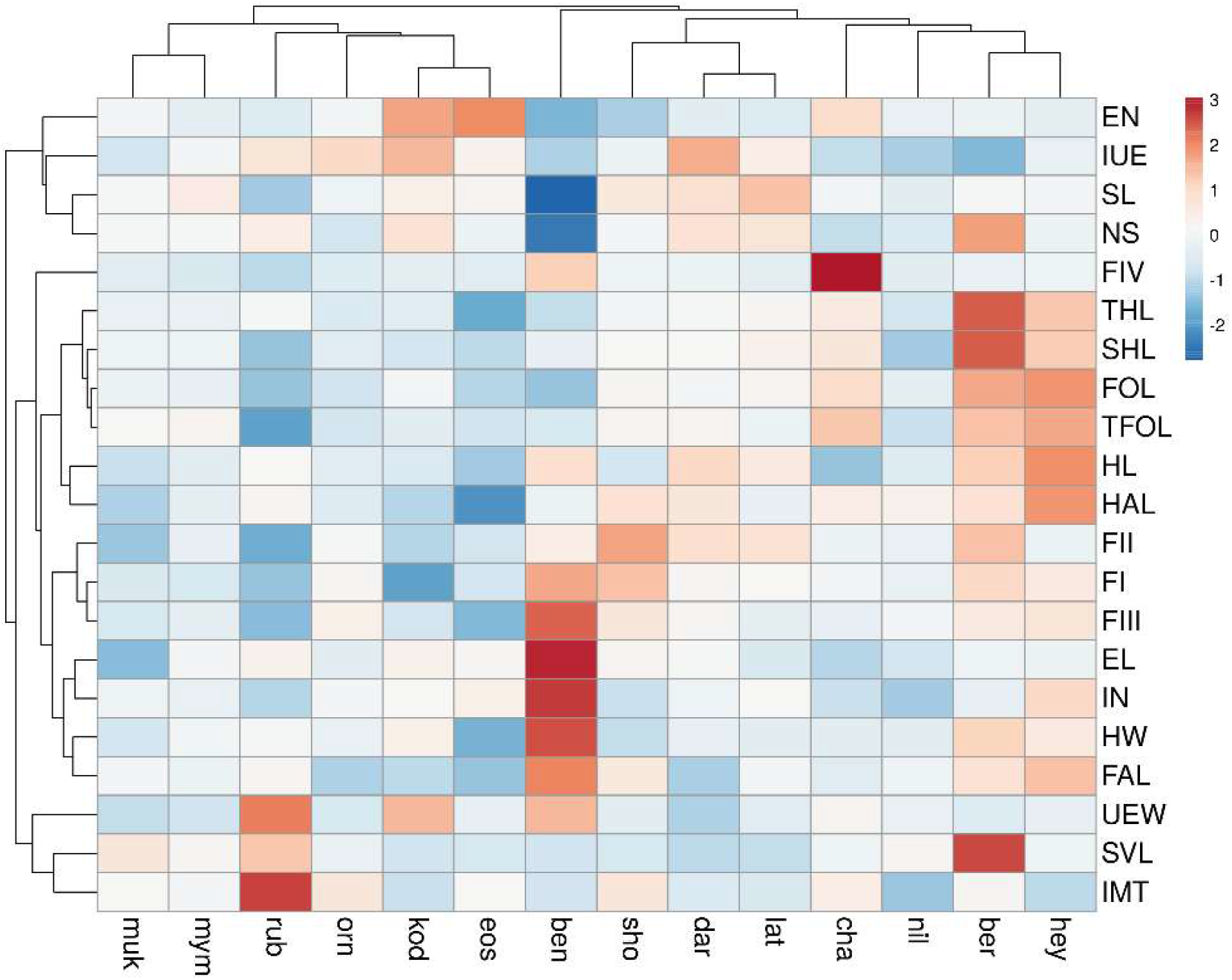
Heat map integrated with phenogram shows morphological variables among 14 species of *Microhyla* using colour code (for abbreviations – species: Table 2, charaters: materials and methods).

### Natural history

*Microhyla bengalensis* is encountered in moist leaf litters of a guava tree in a courtyard of human settlement. There are no water bodies nearby except a drain opposite to the boundary wall. The frog can pass to the drain by a small hole on the boundary wall. So it requires more survey in this region to unravel the mystery of its occurrence in this habitat.

## Discussion

Biju et al (2019) reported 15 species of *Microhyla* from India including *M kodial*. Of these 15 species, occurrence of *M. butleri* and *M. pulchra* is doubtful (Garg et al., 2019). Dinesh et al. (2009) reported occurrence of *M. pulchra* in northeast regions of India. Garg et al. (2019) have the opinion that this could be due to erroneous citing of a report of *Kaloula pulchra* (Dey and Gupta, 2001) and are unable to locate any specimen either in potential museums or during field surveys across India. Lalremsanga et al. (2007) reported *M. butleri* from Mizoram with a snout-vent length 31-34 mm., without any information on sex or vouchers. But previous report on size range of *M. butleri* is 20-25 mm. (male) and 21-26 mm. (female) (Poyarkov et al., 2014). So Garg et al. (2019) commented that report of Lalremsanga et al. (2007) could be a misidentification and is likely to correspond to *M. berdmorei* (male SVL= 33-36 mm.). So number of *Microhyla* species in India should be 13 instead of 15.

Our knowledge on *Microhyla* diversity in India is still far more complete as the species is small *in* size and exploration is restricted mostly in south and north-eastern states. Only three species of *Microhyla* are so far reported from West Bengal namely, *M. ornata*, *M. rubra* (Chanda, 1994) and *M. mymensinghensis* (Biju et al., 2019). Since 2000, 20 new species of *Microhyla* have been discovered globally, mostly in Southeast Asia followed by south Asia (Vineet et al., 2018; Li et al., 2019; Biju et al., 2019; Garg et al., 2019; Matsui et al., 2011; Poyarkov et al., 2014; Gorin et al., 2021).

*M. bengalensis* is close to several small sized species but its incomparable morphological relationship proves its status to be a new species. However, from its distribution (eastern India) and habitat (moist leaf litters), the species is clearly separated from the other known species of *Microhyla* in India. With the discovery of *M. bengalensis* sp. nov., there are now 14 nominated species of *Microhyla* in India and we suspect more species to be revealed by thorough systematic surveys of the poorly explored regions. The highlight of our study is the discovery of a new species in human habitat which is often ignored for amphibian study. Several human activities and development works pose a great threat to destroy the potential habitat of frogs. So engagement of common people to explore the amphibian diversity in local level proves to be effective for conservation of the habitat and the species.

Recently, Gorin et al. (2021) divided *Microhyla-Glyphoglossus* assemblaze into three clades: Clade I- *Microhyla* I (*Microhyla sensu stricto*), Clade II- *Microhyla* II (*Nanohyla* Gen. nov.) and Clade III- *Glyphoglossus*. Of the present 52 species of *Microhyla*, 43 belong to Clade I and nine belong to Clade II. Species of Clade I are widely distributed including India while those of Clade II are restricted to Malayasia and Vietnam. On this basis, the present species belongs to Clade I.

## Acknowledgement

Authors are deeply indebted to Mr. Samriddha Sil for technical help. The first author is grateful to all his colleagues for their constant inspiration.

